# The relevance of accounting for parasympathetic as well as sympathetic arousal in threat conditioning: methodological and clinical considerations

**DOI:** 10.1101/2023.06.06.540498

**Authors:** Lars Jaswetz, Lycia D. de Voogd, Eni S. Becker, Karin Roelofs

## Abstract

Alterations in associative threat learning have been thought to underlie the aetiology and maintenance of anxiety disorders. Recent insights into the facilitatory role of parasympathetic arousal for threat coping have raised the question whether individual differences in parasympathetic versus sympathetic dominance during threat learning may explain the unstable relationship with anxiety vulnerability versus resilience. We applied an established threat-conditioning paradigm in 78 neurotypical individuals and assessed parasympathetic responses (relative bradycardia), as well as sympathetic response patterns (relative tachycardia and increased skin conductance responses -SCR). We observed threat-induced bradycardia as well as tachycardia during associative learning. Additionally, participants not showing conditioned SCR still exhibit significant conditioned threat responses expressed in parasympathetically driven threat bradycardia. Critically, tachycardia, rather than bradycardia, was linked to stronger initial conditioned SCRs and higher trait anxiety. These results suggest individual differences in sympathetic versus parasympathetic dominance may underlie anxiety vulnerability versus resilience.

**Statement of relevance:** Our findings underscore the relevance of assessing the whole spectrum of autonomic nervous system responses to threat. By assessing sympathetic and parasympathetic threat responses, we demonstrate associations with anxiety vulnerability, which could not be unveiled by assessing sympathetic arousal alone. Since alterations in associative threat learning are thought to underlie anxiety-related psychopathology, it is of clinical and methodological relevance to assess threat responses with measures that are sensitive to both parasympathetic and sympathetic arousal. Additionally, we show that individuals that lack sympathetically-driven conditioned SCRs -- often classified as non-learners -- in fact do show a parasympathetically-driven HR threat response (bradycardia). Critically, bradycardia was linked to lower trait anxiety. These results imply a paradigm shift in the field of threat learning, shifting the predominant focus on sympathetic arousal towards the balance between sympathetic and parasympathetic arousal. This could advance insights in the role of threat learning in anxiety vulnerability and resilience.

## Introduction

Deviations in associative threat learning are thought to underlie the development and maintenance of anxiety-related psychopathology (Duits et al., 2015; Marin et al., 2017; Mineka & Oehlberg, 2008; Pittig et al., 2018). Indeed, individual differences in associative learning, assessed by increased sympathetic arousal to conditioned threat, have been linked to vulnerability for anxiety disorders (Sjouwerman et al., 2020). However, these associations do not always replicate (Haaker et al., 2015) and have been largely based on one sympathetic arousal measure, namely skin conductance response (SCR). While heart rate (HR) can capture both sympathetic as well as parasympathetic arousal, previous literature on threat learning has often assessed these responses separately, either classifying sympathetic HR accelerations or tachycardia (Forsyth et al., 2000; Sandin & Chorot, 1989; Wickramasuriya & Faghih, 2020) or parasympathetic HR decelerations, also bradycardia (Ahrens et al., 2016; Bach & Melinscak, 2020; Bradley et al., 2005; Castegnetti et al., 2016; Klorman & Ryan, 1980; Panitz et al., 2015). This one-sided view on conditioned threat responses ignores the scope of the full spectrum. We argue that disregarding individual differences in both response-types during threat learning constitutes a missed opportunity to investigate the conditions under which successful threat learning may be a protective factor or instead signalling vulnerability (Roelofs & Dayan, 2022). We verify whether threat responses can be manifested in both sympathetic and parasympathetic arousal and whether these differences are meaningfully related to trait anxiety. This would imply a paradigm shift in the field of threat learning. Namely, the predominant focus on sympathetic arousal should be shifted towards the balance between sympathetic and parasympathetic arousal. This could advance insights in the role of threat learning in anxiety vulnerability and resilience.

The sympathetic index SCR is by far the most common outcome measure used in human threat learning studies (e.g. Lonsdorf et al., 2017). In such studies, participants learn associations between conditioned stimuli and an aversive outcome such as an electric shock (CS+) or absence of that outcome (CS−). Importantly, one common practice in those studies is that participants that fail to show a discrimination in SCR between a CS+ and a CS− are classified as ‘non-learners’ and are sometimes even excluded from analyses. This procedure has previously been criticized on the grounds that the exclusion criteria remain arbitrary, vary among studies, and may induce a sample bias (see Lonsdorf et al., 2019). Here we go one step further and challenge the classification of ‘non-learners’ itself, asking whether they are genuine non-learners. Solely including SCR as an outcome measure makes it impossible to ascertain whether learning may instead be reflected in parasympathetic arousal, a type of arousal that is as important for coping and can vary independently of sympathetic arousal (Löw et al., 2008; Roelofs & Dayan, 2022; Van Diest et al., 2009).

Acute threat activates both the parasympathetic and sympathetic branch of the autonomic nervous system (ANS). The relative balance of activity in these systems is associated with distinct response patterns including freezing and fight-flight reactions, respectively (Eilam, 2005; Fanselow, 1994; Roelofs, 2017; Trott et al., 2022). Freezing is a parasympathetically dominant state accompanied by motor inhibition and HR deceleration (i.e. a bradycardic response; Hermans et al., 2013; Noordewier et al., 2020; Roelofs et al., 2010; Walker & Carrive, 2003), while fight-or-flight reactions reflect a sympathetically dominant state accompanied by a HR acceleration (i.e. a tachycardic response; Hagenaars, Oitzl, et al., 2014; Hagenaars, Roelofs, et al., 2014; Iwata & LeDoux, 1988; Keay & Bandler, 2001; Walker & Carrive, 2003; Waxenbaum et al., 2021). Which branch is more dominant can depend on situational factors, such as threat proximity (D. C. Blanchard et al., 2011; Löw et al., 2008; Mobbs, 2018; Mobbs et al., 2020), escape possibilities (R. J. Blanchard & Blanchard, 1968; Qi et al., 2018), or whether a later action is required (Gladwin et al., 2016; Roelofs & Dayan, 2022).

Interestingly, an observation not frequently highlighted in the literature is that in addition to situational factors, there are also inter-individual differences in which ANS branch is more dominant in response to acute threat. Namely, some individuals tend to show relative HR decelerations (i.e. bradycardia) in response to threat, while others have the tendency to show HR accelerations (i.e. tachycardia) to the same threat context (Cohen & Randall, 1984; de Echegaray & Moratti, 2021; Hodes et al., 1985; Moratti & Keil, 2005; Sevenster et al., 2015; van Ast et al., 2022; Van Diest et al., 2009). It is not yet clear, what underlies these different response patterns. Parasympathetic dominance has been associated with enhanced perception and decision-making under threat (de Voogd et al., 2022; Hashemi et al., 2019; Livermore et al., 2021; Lojowska et al., 2015, 2018; Ly et al., 2014; Rösler & Gamer, 2019). Additionally, ANS responses are often altered in individuals with anxiety-related psychopathology (Stone et al., 2020; Vinkers et al., 2021), suggesting that parasympathetically dominant threat-responding constitutes an important factor to consider when studying individual differences in anxiety vulnerability and resilience. Here we explore the possibility that inter-individual differences in relative bradycardia and tachycardia responses reflect different associative learning processes that in turn may be related to vulnerability and resilience factors underlying anxiety-related psychopathology.

Indeed, there is evidence that ANS activation and response tendencies to threat are related to the development and maintenance of psychopathology. For instance, shorter freezing duration or no freezing at all in response to an acute threat during early childhood is predictive of later internalizing symptom development (Held et al., 2022; Niermann et al., 2017). Analogous to that, hyperactivity of the sympathetic nervous system has been linked to trauma-related symptoms. For instance, patients with post-traumatic stress disorder (PTSD) and/or anxiety disorders consistently show dysregulated ANS activity, such as tachycardia, heightened skin conductance, and blood pressure (R. E. Bernstein et al., 2013; Blechert et al., 2007; Brawman-Mintzer & Lydiard, 1997; Lipov, 2013; Morris & Rao, 2013; Pole, 2007). Moreover, PTSD patients tend to show tachycardia to threatening stimuli compared to controls who typically show bradycardia (Adenauer et al., 2010; Fragkaki et al., 2017). Finally, pre-deployment parasympathetic dominance in resting HR variability has been linked to increased stress-resilience in marines (Minassian et al., 2015), while higher resting HR after traumatic injury predicted subsequent PTSD development in children (Bryant et al., 2007).

Together these finding suggest that it is relevant to not only assess sympathetic but also parasympathetic arousal when evaluating threat learning. Particularly because parasympathetic dominant responses to threat may be linked to resilience and to reduced manifestation of vulnerability factors underlying anxiety-related psychopathology, such as trait anxiety (Knowles & Olatunji, 2020). Investigating these relations, is not only relevant to advance our understanding of threat learning, but the findings could also impact methodological considerations in threat conditioning paradigms, where exclusion criteria are often hinged on sympathetic (i.e. skin conductance, see Bach et al., 2023; Lonsdorf et al., 2017). Previous work has already indicated that exclusion practices based on non-learning could greatly affect the conclusions drawn from such studies (Lonsdorf et al., 2019). Here we explore an additional problem, namely the possibility that those non-learners based on SCRs may in fact show learned responses on HR in terms of bradycardia, a pattern that may be relevant for optimal coping and reduced anxiety (Roelofs & Dayan, 2022). Therefore, measurement choices may not be arbitrary but may reflect qualitative different processes.

To test our hypotheses, we used an established differential delay threat conditioning paradigm (Jaswetz et al., 2022), during which both HR and SCR were measured. Our sample included a relatively large number of participants (*N*=78) allowing us to explore relations with trait anxiety. As preregistered, based on previous findings (de Echegaray & Moratti, 2021; Sevenster et al., 2015) our first aim was to verify that associative threat reactions can be manifested in both bradycardia as well as tachycardia. Second, we additionally assessed whether those HR response patterns were also correlated to sympathetically driven SCR, and to various classes of SCR-based “non-learners”. Thirdly, we tested the possibility that the magnitude of both threat-induced HR bradycardia as well as tachycardia may be related to trait anxiety, a marker that has been linked to vulnerability to develop anxiety-related psychopathology.

## Method

This study was preregistered on the open science framework (Link: https://osf.io/48eqs). As this study was part of a previous project on threat memory reconsolidation (Jaswetz et al., 2022), the preregistration was written after visual inspection of the data, but before conducting any statistical analyses of the data relevant to this study.

### Participants

We recruited healthy individuals through the online recruitment system of the Radboud University. Inclusion criteria were: above the age of 18, with normal or corrected to normal vision, no acute mental disorder, no skin disease that would prohibit the use of electrodes, and no history of brain trauma or brain surgery. In total, 78 individuals (49 females, 29 males, 18–60 years [*M* = 24.73, *SD* = 7.05]) completed the entire study. There was one individual who terminated the experiment early. All participants provided informed consent and received €16 as a compensation for their participation.

### Procedure

Participants were tested in a differential delay threat conditioning paradigm (Jaswetz et al., 2022). Participants came to the lab and filled in an informed consent for the entire study, as well as a screening list, and two questionnaires (see below). Next, participants were instructed to wash their hands to clean off any soap or disinfectant and, in case their hands were cold, to warm up their hands. Next, participants completed the shock workup procedure to calibrate the intensity of the electric shock (see below). Afterwards, participants were subjected to the acquisition phase of threat conditioning paradigm. Finally, participants filled in a five-point rating scale concerning their shock expectancy and subjective feelings regarding the likeability of each stimulus. The data presented here was collected as part of a project on memory reconsolidation and included two more testing sessions (Jaswetz et al., 2022) not included here.

### Materials

#### Conditioned stimuli

The conditioned stimuli (CS) consisted of three rectangles in the colours blue, green, and yellow. Two of these stimuli were paired with an electrical shock. Additionally, one stimulus was never paired with a shock. Assignment of colours to the different CS types was counterbalanced across participants. For this study, we averaged responses across the two CS+ trials.

#### Unconditioned stimulus

The unconditioned stimulus (UCS) was an electrical shock that was delivered via two Ag/AgCl electrodes attached to the distal phalanges of the second and third finger of the left hand. The shock was delivered via a MAXTENSE 2000 (Bio-Protech) electrical stimulation machine, with a frequency of 140 Hz and a duration of 200 ms. The shock intensity was set during a standardised shock workup procedure (de Voogd et al., 2016; Klumpers et al., 2010) and remained the same throughout the whole experiment. In this procedure all participants received five shocks. After each shock, participants subjectively rated the experienced unpleasantness on a scale ranging from 1 (not painful at all) to 5 (very painful), based on which the subsequent shock was adjusted, in order to arrive at a shock intensity that was unpleasant, but not painful. The intensity varied in 10 intensity steps between 0 and 40 V/ 0–80 mA. The average intensity step was M = 4.48 (SD = 1.87).

#### Differential delay threat conditioning paradigm

Participants were instructed to discover the relationship between the CSs and the UCS. The acquisition phase consisted of 48 trials in total (32 CS+ trials, 16 CS− trials). The stimuli were presented in a pseudo randomised order, with no more than three stimuli of the same type being presented in succession. Each CS (4 s duration) was followed by an inter-trial interval (ITI) during which a fixation cross was shown (jittered 6–10 s, *M* = 8 s duration). For the CS+ trials, the shock (200 ms duration) was delivered at 3.8 s after stimulus onset. Reinforcement rate was set at 37.5% for each CS+, meaning that 12 out of 32 CS+ presentations were paired with a shock. The first presentation of each CS+ was always paired with a shock to facilitate immediate and equal learning for both CS+ types. The remainder of the shocks were pseudo randomly distributed across the first and second half of the acquisition phase. This was done to ensure that the shocks were spread evenly across the whole acquisition phase.

#### Physiological measures

Electrodermal activity (EDA) and HR were measured throughout the experiment (5000 Hz). EDA was measured via two Ag/AgCl electrodes attached to distal phalanges of the first and second finger of the right hand. HR was measured via a pulse sensor attached to the third finger of the right hand. Additionally, a woollen gauntlet was pulled over the participant’s right hand to keep the hand warm during the experiment.

#### Questionnaires

Trait anxiety was measured with the trait subscale of the State Trait Anxiety Inventory (STAI-T; Spielberger, 1983) and childhood adversity with the Childhood Trauma Questionnaire (CTQ; D. P. Bernstein et al., 1998). Since the CTQ scores were right-skewed and leptokurtic (skewness = 1.48, kurtosis = 5.06) we were unable to use CTQ scores as an inter-individual difference measure (see Blanca et al., 2013). The STAI scores were not skewed and only slightly platykurtic (skewness = 0.38, kurtosis = −0.31). In our sample, the STAI showed a Cronbachs alpha of *α* = 0.89, indicating good reliability (Tavakol & Dennick, 2011). The STAI sum scores were computed by first inversing all item scores of mirrored items and then adding all item scores.

### Preprocessing

The raw physiological data were first preprocessed using inhouse software (brainampconverter: https://github.com/can-lab/brainampconverter). The downsampled (100 Hz) HR data were then scored using inhouse software (hera: https://github.com/can-lab/hera) for peak detection (with additional manual supervision). The resulting inter-beat interval (IBI) time course data was then exported to R (R Core Team, 2020) and transformed into beats per minute (BPM). For each trial, a baseline window (1 second before trial onset up to trial onset; de Voogd et al., 2022) was taken and subtracted from the trial window (1 – 3.8s after trial onset), resulting in a baseline corrected average BPM per trial. Grouping participants into threat-induced bradycardia and tachycardia groups was done based on the average bradycardia and tachycardia response per participant. These were determined by subtracting average BPM to the CS+ minus average BPM to the CS−, where a negative value indicates threat bradycardia, and a positive value indicates threat tachycardia.

After down sampling (200 Hz), skin conductance responses (SCR) data were scored with additional manual supervision using Autonomate (Green et al., 2014) implemented in Matlab (*MATLAB*, 2018). Here, the amplitude of a rise in SCR was scored. The rise had to start between 0.5 s after stimulus onset and 0.5 s after stimulus offset, with a minimum rise time of 0.5 s and a maximum rise time of 5 s after response onset. The SCRs from the acquisition phase were normalised to the average shock SCRs during the acquisition phase and square root transformed. All reinforced trials were excluded from the analyses involving SCR but were included in analyses involving HR as the shock was delivered outside the analysis window.

### Data-analyses

The data were analysed with Bayesian Mixed Effects Models in R (R Core Team, 2020) using the *brms* package (Bürkner, 2018).

To assess successful conditioned HR responses, we first ran a model with Baseline Corrected BPM, for each trial, as the dependent variable and CS Type (CS+, CS−) and Trial Number (1-16) as fixed effects, with a random slope for both fixed effects over a random intercept for Participant ID. We then, to investigate the association between HR and our other variables of interest, split up the group the based on whether participants expressed a threat-induced bradycardic versus tachycardiac response pattern (similar to de Echegaray & Moratti, 2021).

Next, to assess the relation between parasympathetically and sympathetically dominant response patterns, we investigated the relationship between HR and SCR. We ran a model with SCR as the dependent variable and CS Type (CS+, CS−) and in addition HR Groups (threat tachycardia and bradycardia group) as fixed effects and with a random slope for CS Type over a random intercept for Participant ID. We then followed this up by including Trial Number as a fixed effect in that model (not preregistered) to assess time effects as well. We ran follow-up models, in case of significant main or interaction effects. We deviated from the preregistration because we proposed to use the HR as the dependent variable. The advantage of the current analysis is that it is more intuitive and informative for the understanding of the relationship between HR and SCR to make SCR the dependent variable which was our intention of the analysis.

Next, our main analysis of interest involved to assess the relation between HR response patterns and Trait Anxiety. Therefore, we ran a model with HR as the dependent variable and CS Type (CS+, CS−) and Trait Anxiety as an additional fixed effect, with a random slope for CS Type over a random intercept for Participant ID. Because there is also the possibility that the magnitude of both HR bradycardia and tachycardia (regardless of the direction of HR change) is linked to Trait Anxiety, we ran an additional model (as preregistered) with the absolute difference score between the CS+ and CS− on HR as the dependent variable and Trait Anxiety as the fixed effect, with a random intercept for Participant ID.

Finally, the results from the 2^nd^ model gave rise to a further (not preregistered) analysis of whether classifications of SCR non-learners also followed non-learning in HR. To better understand the consequences of the results of the first step we first computed SCR difference scores between the CS+ and CS−. Participants scoring above cut-off scores 0.2, 0.1, and 0 normalised and square rooted µS were labelled as learners (see Lonsdorf et al., 2019 for a similar approach). Participants scoring below the respective cut-offs were labelled as non-learners. We first a strict learning criterion (CS+ - CS− above 0.2 normalised and square rooted µS), meaning that 76% (59 out of 78) individuals did not fulfil the learning criterion. We assessed whether this group showed successful conditioned HR responses. As this was a large group, we applied two more lenient learning criteria (CS+ - CS− above 0.1 normalised and square rooted µS, and CS+ - CS− above 0 normalised and square rooted µS). Now, only 56% (44 out of 78) and 21% (16 out of 78) individuals did not fulfil these learning criteria. To clarify, individuals included in the lenient cut-offs were also included in the stricter cut-offs, therefore these analyses were not independent. We then analysed whether there was significant differentiation on HR responses within the SCR non-learner groups by running a simple brms model per cut-off score with HR as the outcome variable and CS Type (CS+, CS−) as the fixed effect, with a random slope for CS Type over a random intercept for Participant ID.

Lastly, we also assessed whether the absolute magnitude, rather than the direction (i.e. threat bradycardia/tachycardia) of the HR threat response was associated with individual differences in sympathetic arousal and Trait Anxiety. To that end, we ran models with the absolute difference between the CS+ and CS− in BPM per participant as the dependent variable and SCR as well as Trait Anxiety as the fixed effects, respectively. We preregistered additional exploratory analyses but opted not to report them here.

Since Bayesian analyses do not yield p-values but instead work with 95% confidence intervals (CI), effects were considered “significant” in the traditional sense when the 95% confidence interval of the posterior distribution did not include zero. In addition to the confidence interval, we also report the estimate. The estimate is the mean of the posterior distribution, which is the probability distribution of the parameters conditional on the data. We opted not to use Bayes Factors as the test statistic since we did not have informative priors. We tried to use a maximal model approach for our random effects structure (see Barr, 2013), however, we sometimes had to forego random slopes due to convergence issues.

## Results

### Threat-induced bradycardia as well as tachycardia

First, we tested whether overall differential threat learning was expressed in HR responses. In line with previous work (Castegnetti et al., 2016; Klorman & Ryan, 1980), our results showed that the CS+ elicited overall significantly lower BPM than the CS− (estimate = −0.84, 95% CI[−1.38; −0.31]) and that the overall BPM significantly rose across trials (estimate = 0.11, 95% CI[0.04; 0.19]), but that the change in BPM across trials did not differ between CS+ and CS− (estimate = −0.02, 95% CI[−0.11; 0.06]). In line with expectations, we observed both patterns of threat-induced HR bradycardia (N=49) and HR tachycardia (N=29), when numerically dividing participants based on their BPM difference score between the CS+ and CS− (i.e. de Echegaray & Moratti, 2021 – see **Figure 1**).

**Figure 1.**
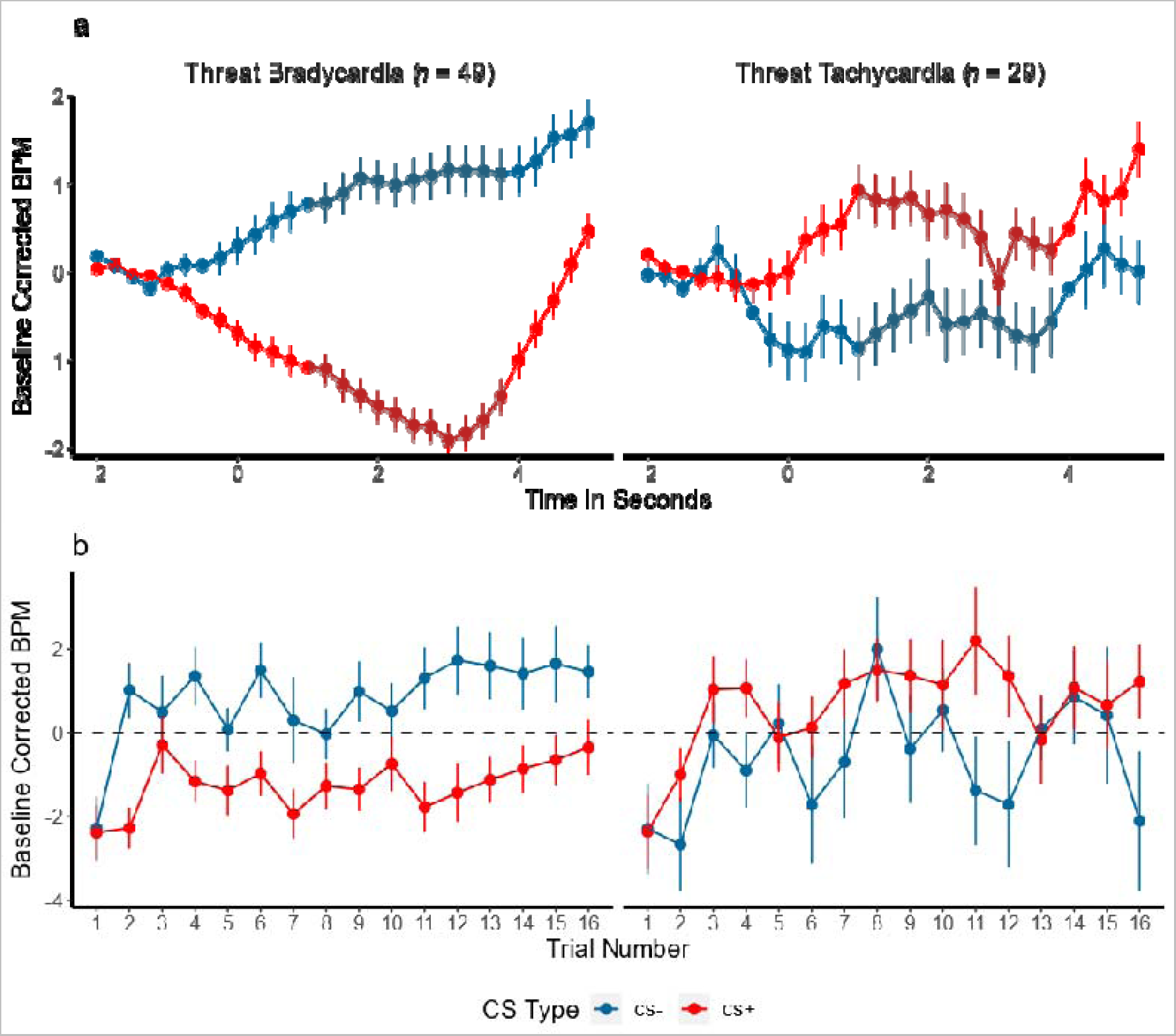
HR responses in threat-conditioned bradycardia and tachycardia response groups. a) Average time course of the HR across a trial, split by CS Type and threat-induced bradycardia and tachycardia groups. b) HR across all trials, split by CS Type and threat-induced bradycardia and tachycardia groups.

### Threat-induced tachycardia is associated with heightened sympathetic arousal

Next, we tested whether sympathetically dominant response patterns of HR in the tachycardia responders are related to the sympathetic index SCR. As commonly observed, we found a significant time effect (Jaswetz et al., 2022), showing that SCRs decreased over successive trials, possibly due to habituation. Across the entire time course, there was no significant difference in average conditioned SCR responses (CS+ vs CS−) between the HR tachycardia and bradycardia group (estimate = 0.03, 95% CI[−3.02; 3.11]). However, the change in conditioned SCR responses (CS+ vs CS−) over time was significantly different between the groups (estimate = −0.01, 95% CI[−0.02; −0.01]). Follow-up analyses to explain this effect, indicated that the differences in the change in SCRs across time between HR Groups (threat bradycardia and threat tachycardia) was significantly different for the CS+ (estimate = 0.01, 95% CI[0.001; 0.01]), but not for the CS− (estimate = −0.00, 95% CI[−0.00; 0.00]). The difference in CS+ SCRs between the groups was more pronounced during the first compared to the second half of the trials (estimate = −0.02, 95% CI[−0.03; −0.01]) indicating the groups to mainly differ in SCRs during early learning.

We also assessed whether the absolute difference in HR between the CS+ and CS− (i.e. the magnitude of the threat response, regardless of direction) was related to SCR. Here, we analysed the absolute BPM difference between the CS+ and CS− across SCR. The results show that there is no significant relation between the absolute HR difference and SCR (estimate = −1.72, 95 CI [−4.36, 0.92]).

Thus, there was a difference between the bradycardia and tachycardia HR Groups in conditioned SCRs (characterized by a steeper slope over trials) that was mainly driven by the CS+ in the first half of the trials. Together, these results indicate that individual differences in bradycardia and tachycardia response patterns are also characterized by differences in skin conductance response patterns, a main measure of sympathetic arousal (See **Figure 2**). Importantly, the sympathetic measure SCR was not associated with the magnitude of the HR responses, but rather the direction of the threat related HR response (threat tachycardia vs. threat bradycardia).

**Figure 2.**
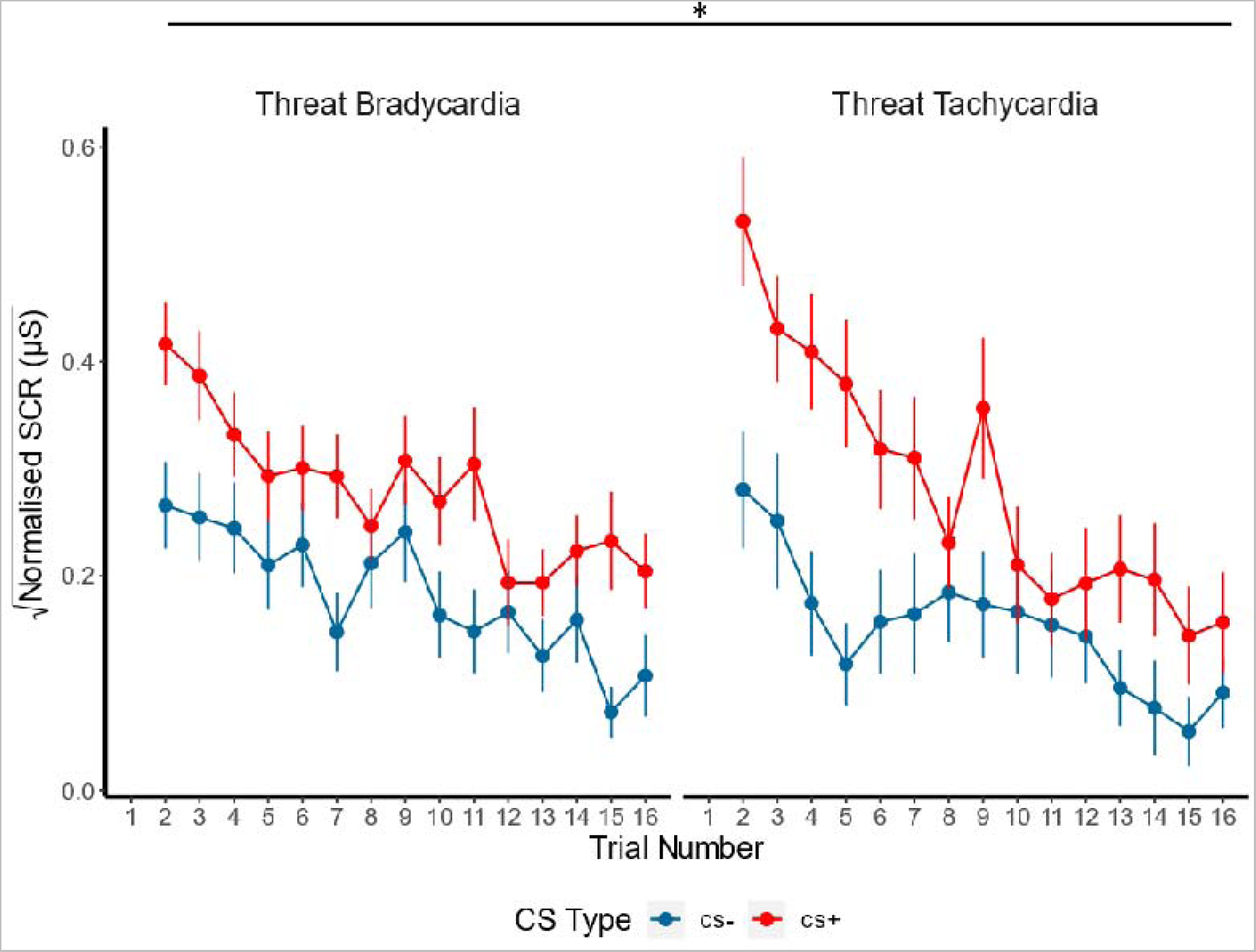
SCR across all trials split by CS Type and threat-induced HR bradycardia and tachycardia groups. The Figure illustrates stronger conditioned skin conductance responses (SCR) for the CS+ trials in the tachycardia vs. bradycardia group, at the start of the experiment in particular.

### SCR non-learners show learning in terms of parasympathetic arousal

The next analysis was inspired by the notion that participants that fail to show a differential SCR response (CS+ > CS−) are sometimes classified as “non-learners” and -depending on the context of the study-excluded from further analyses. Here we assess whether these individuals may show differential HR responses (i.e. on an index that is sensitive to parasympathetic arousal) regardless. Here, we used three increasingly liberal (with respect to including only learners in the sample) cut-off values to assess differentiation between the CS+ and CS− on HR responses within the excluded group e.g. SCR non-learners. For the most conservative cut-off value (CS+ minus CS− above 0.2 normalised and square rooted µS), which included 59 participants as “SCR non-learners”, there was a significant difference between the CS+ and CS− on HR responses (estimate = −0.91, 95% CI[−1.52; −0.29]). Next, with a more lenient cut-off (CS+ minus CS− above 0.2 normalised and square rooted µS, *n* = 44), we also found a significant difference between the CS+ and CS− on HR responses (estimate = −1.08, 95% CI[−1.79; −0.37]). Lastly, for the most lenient cut-off (CS+ minus CS− above 0.2 normalised and square rooted µS, *n* = 16), which represents the smallest group of “SCR non-learners”, we still found a marginally significant difference between the CS+ and CS− on HR responses (estimate = −0.96, 95% CI[−1.95; 0.04], 90% CI [−1.78, −0.14]). For an overview of these results, See **Figure 3**). These results indicate that SCR non-learners, i.e. individuals do not show a differential SCR response, do seem to show differential HR responses that on average are manifested as bradycardia. These findings corroborate the findings from the previous section by further providing evidence that bradycardia response patterns are accompanied by attenuated differential SCR.

**Figure 3.**
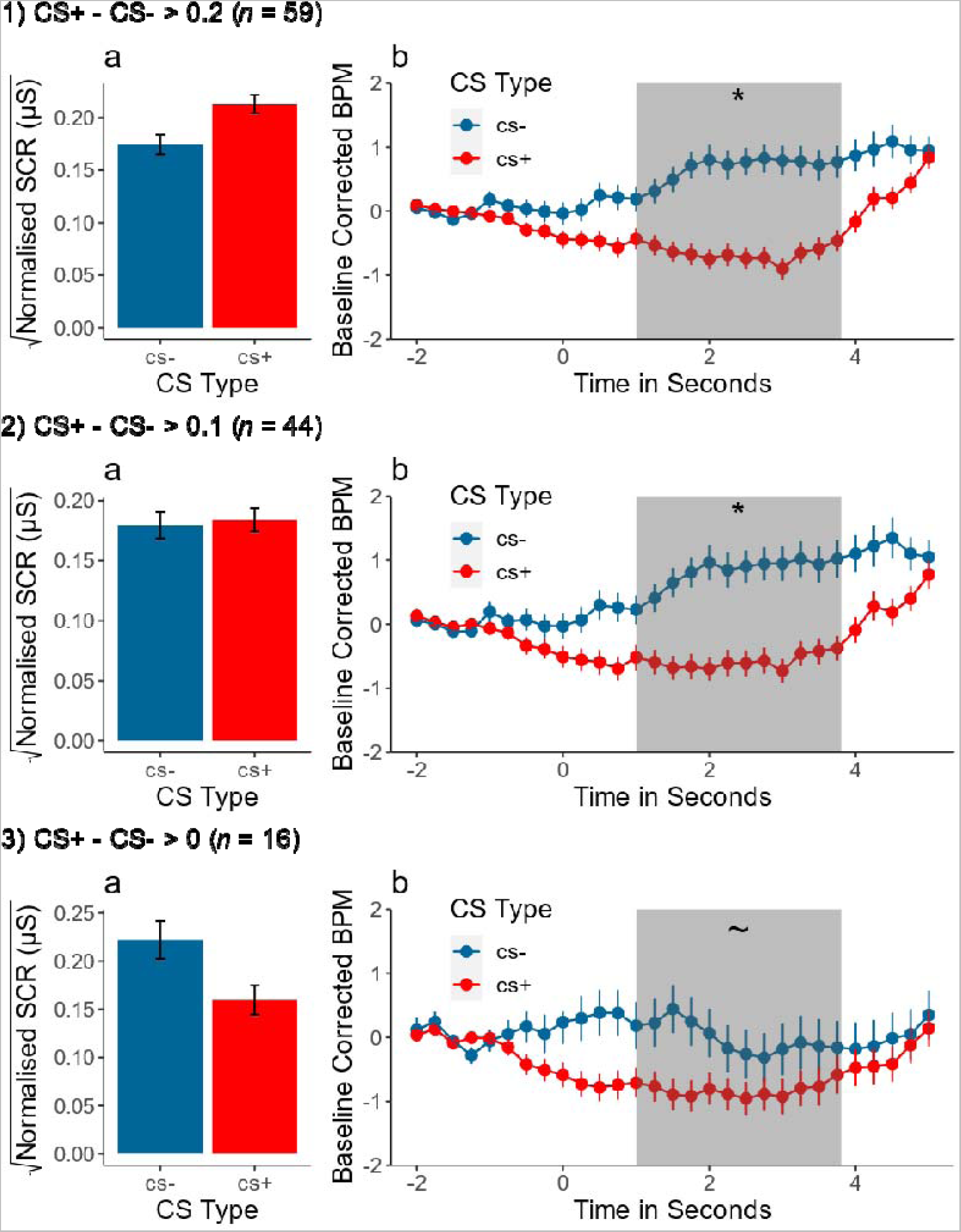
Heart rate responses in SCR-based ‘non-learners’ indicate learning in terms of parasympathetic arousal instead of sympathetic arousal. a) SCR to the CS+ and CS− within the non-learner group (i.e. difference between CS+ and CS− scoring below the respective cut-off value. b) HR response to the average CS+ and CS− trial within the SCR non-learner group. Shaded area represents the analysis window. 1), 2), and 3) represent the respective SCR cut-off values. The Figure illustrates that sympathetic ‘non-learners’ according to varying previously defined SCR-criteria do show learning on a parasympathetic arousal measure: HR deceleration.

### Threat-induced HR bradycardia responders have relatively low trait anxiety

After having established that associative threat learning can be reflected in both sympathetic as well as parasympathetic arousal, we verify the clinical relevance of both types of response patterns. In light of previous conflicting results for predictability of trait anxiety by threat conditioned SCR responses, we deemed it particularly relevant to assess the relationship between threat-induced bradycardia and tachycardia responses and trait anxiety. First, we ran a group-based model assessing whether threat-induced bradycardia and tachycardia HR Groups differed in their levels of Trait Anxiety. The results showed that the groups differed significantly, with the bradycardia responders showing lower Trait Anxiety scores as compared to the tachycardia responders (estimate = −3.85, 95% CI[−5.81, −1.89]). See **Figure 4b**. Next, we ran a model, assessing whether HR responses to the CS+ and CS− were related to trait anxiety, across individuals rather than between groups. Indeed, overall lower HR response patterns was related to trait anxiety. (estimate = −0.05, 95% CI[−0.09; −0.01]). Importantly, there was a significant interaction effect, indicating that the relation between trait anxiety and HR differs between the CS+ and CS− (estimate = 0.10, 95% CI[0.04; 0.16]). With increasing Trait Anxiety, the CS− elicited lower HR responses while the CS+ elicited higher HR responses (for a visual representation of the results, see **Figure 4a**). Finally, we also assessed whether Trait Anxiety was linked to altered differentiation between the CS+ and CS− in terms of HR responses *regardless of the direction*. To that end, we analysed the *absolute* difference in HR responses between the CS+ and the CS−. The results showed a significant main effect of Trait Anxiety (estimate = −0.05, 95% CI[−0.09; −0.01]), indicating that also the absolute difference in HR responses between CS+ and CS− decreased with increasing trait anxiety (see **Figure 4c**). Together these findings suggest that tachycardic response patterns constitute an important marker for higher trait anxiety and that with increasing anxiety, the discriminative ability (between CS+ and CS−) is impaired in terms of HR responses. This indicates that HR-besides providing an often ignored but important index of parasympathetic arousal in associative threat learning-can reflect differential response patterns that are meaningfully related to discriminative learning and trait anxiety.

**Figure 4.**
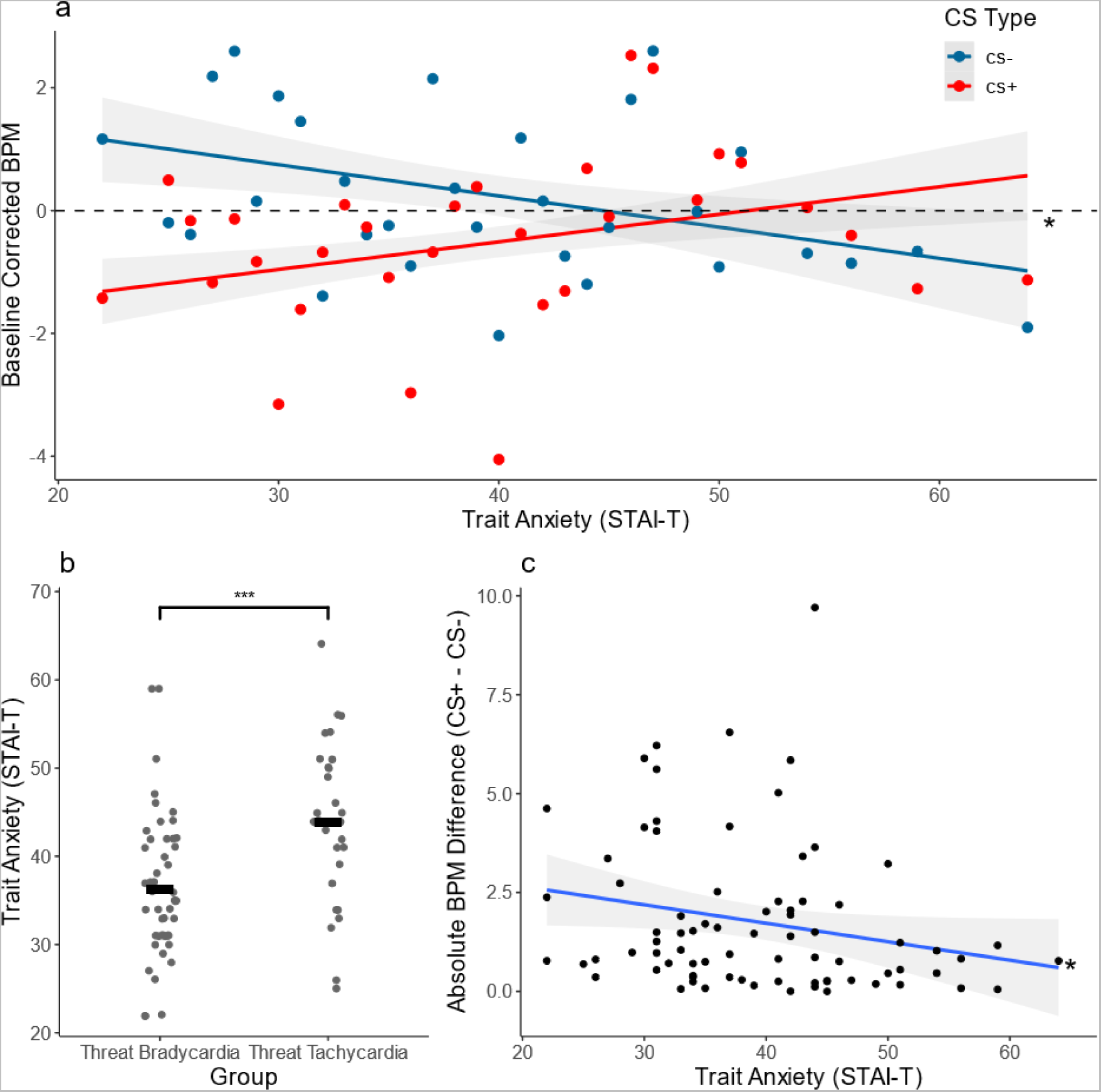
Threat conditioned HR bradycardia is linked to low trait anxiety. a) Trait Anxiety levels of bradycardia and tachycardia HR Groups. Black lines indicate mean values per group. b) Absolute BPM differences (baseline corrected) between CS+ and CS− are correlated to Trait Anxiety scores. c) Relation between HR (Baseline corrected BPM) and Trait Anxiety scores split by CS Type. This panel illustrates, that participants with stronger CS Type discrimination (hence stronger threat-induced bradycardia) have lower Trait Anxiety scores.

## Discussion

The aim of this study was to test whether the consideration of individual differences in sympathetic versus parasympathetic dominance during threat learning may shed light on the complex relationship with anxiety vulnerability and resilience. First, we verified that threat conditioning can be reflected in bradycardia as well as tachycardia types of response patterns. Critically, and supporting the relevance to consider parasympathetic as well as sympathetic arousal during threat conditioning, our results indicated that tachycardia responders showed stronger initial conditioned SCRs (indicated by a steeper slope during early learning) and higher trait anxiety compared to bradycardia responders. Next, we tested the relevance of these findings for current common practice in the extensive field of threat learning where ‘non-learners’ are typically exclusively based on sympathetic arousal indices (SCR) and are sometimes even excluded from analyses. Using frequently applied ‘SCR non-learner’ criteria (Kredlow et al., 2017; Lonsdorf et al., 2019), we showed that individuals previously classified as ‘non-learners’ were in fact showing successful learning in terms of parasympathetic arousal. Methodological as well as clinical implications of these findings will be discussed.

The finding that both parasympathetically dominant (indicated by bradycardia) as well as sympathetically dominant (tachycardia) threat responses can be observed during associative threat learning is in line with earlier observations in both human and animal literature (de Echegaray & Moratti, 2021;; Hodes et al., 1985; Plunkett, 1979; Sevenster et al., 2015; for a review see Haaker et al., 2019), where such patterns were found in individuals and animal subjects confronted with various forms of (learned) threat. We extend these findings by showing that tachycardia is related to altered SCR patterns, namely stronger initial conditioned responses. Moreover, we show that distinguishing HR response types is meaningful both in terms of SCR and CS discrimination and in terms of trait anxiety levels. These findings call for a paradigm shift in the field of threat learning, where in recent years, sympathetic responses have been favoured over HR measures (for an overview, see Lonsdorf et al., 2017), and direct comparisons between sympathetic and parasympathetic dominance in threat-response patterns have been lacking. We feel such shift is needed for several reasons. First, there is a growing body of literature indicating that in response to anticipation of acute threat, sympathetic and parasympathetic ANS are both activated, and that parasympathetic arousal is in fact dominant in most cases (Roelofs, 2017; Roelofs & Dayan, 2022; van Ast et al., 2022). Parasympathetic dominance is featured by bradycardia as well as tonic immobility and often jointly referred to as freezing (Gladwin et al., 2016; Hagenaars, Roelofs, et al., 2014; Klaassen et al., 2021; Niermann et al., 2018). There is increasing evidence that threat-anticipatory freezing can occur during associative threat learning (Battaglia et al., 2022; Castegnetti et al., 2016; Klorman & Ryan, 1980; Szeska et al., 2021) and is relevant for active threat coping (Gladwin et al., 2016; Hashemi et al., 2019; Roelofs, 2017). Second, we see a meaningful relation between the sympathetic measure SCR that is most used in threat learning studies and HR responses, but the relation is not a one-to-one relation. Hence SCR cannot replace the HR responses, which are predominantly parasympathetically controlled. In our study, threat-induced tachycardia responders did show stronger sympathetically driven conditioned SCRs compared to threat-induced HR decelerators, but that was mainly during early learning, indicating that both measures have unique value. Third, when we classified individuals as SCR “non-learners” according to several increasingly conservative exclusion criteria (for an overview of such practices, see Lonsdorf et al., 2019), we show that SCR non-learners still showed discriminatory threat-induced HR deceleration responses. Thus, SCR “non-learners” should not be considered per se as non-learners, but could be characterized as individuals that show a learned parasympathetic dominant response instead. Fourth, parasympathetic and sympathetic responders showed differential trait anxiety levels, with parasympathetic responders showing a profile in line with relative resilience. Together, these findings suggest it is important for future studies to take both sympathetic and parasympathetic response patterns into account when assessing individual differences in associative learned threat responses and argue for a broader view on what constitutes threat-induced arousal, encompassing both branches of the autonomic nervous system.

Particularly, the finding that sympathetic or parasympathetic dominance during threat were differentially linked to individual differences in trait anxiety deserves additional discussion. Threat-induced bradycardia was associated with lower levels of trait anxiety whereas a pattern of threat-induced tachycardia was associated with higher levels of trait anxiety. Although never jointly studied in threat conditioning per see, these results are broadly in line with earlier research on ANS reactivity in anxious psychopathology. For instance, previous studies showed that indices of anxious psychopathology were related to less parasympathetic and more sympathetic ANS activity (Adenauer et al., 2010; R. E. Bernstein et al., 2013; Brawman-Mintzer & Lydiard, 1997; Held et al., 2022; Lipov, 2013; Morris & Rao, 2013; Niermann et al., 2017; Pole, 2007). In the context of threat, one explanation for these differences in ANS responses could be that anxious individuals tend to mobilise for avoidance (see Hodes et al., 1985), which is then accompanied by sympathetic dominance (i.e. flight or fight reactions). Hamm and colleagues (1993) discuss that cardiac acceleration in response to unpleasant stimuli (i.e. threat-induced tachycardia) only occur in phobic (i.e. highly anxious) individuals, which is interpreted as serving defensive action preparation. Indeed, previous literature on avoidance behaviour under threat shows that anxious psychopathology is associated with more pronounced avoidance tendencies (Hulsman et al., 2021; Pittig et al., 2021; Pittig & Scherbaum, 2020). Analogous to that, mobilising for defensive action has been associated with sympathetic arousal (Van Diest et al., 2009). In studies with unavoidable and avoidable threat, tachycardia responses were seen when participants took action to avoid threat, whereas bradycardia responses were seen when participants encountered an unavoidable threat (Löw et al., 2015; Wendt et al., 2017). It is therefore possible that highly anxious individuals under threat tend to prepare for action (i.e. avoidance), even when no action can be taken. Therefore, parasympathetically dominant HR bradycardia as well as sympathetically dominant HR tachycardia can both be seen as ANS reactions to a learned threat, albeit tachycardia may contain an impulsive action preparation component that is related to fearful avoidance tendencies, even when avoidance might not be optimal.

The finding that differences in HR responses (i.e. the magnitude of the threat response regardless of direction) between the threatening and the safe stimuli decreased with increasing trait anxiety might be indicative of threat generalisation. Threat generalisation is typically characterised by the presence of threat responses to innocuous stimuli (such as the safe stimulus in this study), that resemble responses evoked by threatening stimuli (for a review, see Dymond et al., 2015). In line with this notion, a recent meta-analysis has shown that indices of anxious psychopathology are linked to fear generalisation in threat learning (Sep et al., 2019). Therefore, indices of anxious psychopathology could be linked to ANS responses to threat which not only resemble avoidance reactions, but also generalise to innocuous stimuli.

The above discussion is largely centred around associative learning where parasympathetic dominance may reflect an adaptive response pattern at post-encounter threat, that helps action preparation, upregulated sensory processing, and decision making (de Voogd et al., 2022; Gladwin et al., 2016; Roelofs & Dayan, 2022) and is linked to low anxiety (Held et al., 2022; Niermann et al., 2017). Studies on non-associative acute threat provide a somewhat distinct picture in terms of individual differences in trait anxiety. For instance, some studies assessing HR responses to aversive pictures found that higher indices of anxiety were related to stronger bradycardia responses instead of tachycardia responses (Hagenaars et al., 2012; Noordewier et al., 2020; Roelofs et al., 2010), while other studies on non-associative acute threat did not find a discernible link between trait anxiety and HR responses (Hashemi et al., 2021; Rösler & Gamer, 2019; Sege et al., 2018). This discrepancy could stem from design-specific factors that differ across the mentioned studies. Where associative threat induces a threat-anticipatory state marked by action preparation and heightened alertness in response to a CS (post-encounter threat), non-associative threat responses may as well reflect reaction to the UCS (circa strike), which is not marked by attentive immobility and bradycardia. Indeed, differences in situational characteristics and demands may play a role. Several studies showed that HR responses differed depending on whether an action could be taken to avoid/escape a threatening situation (Gladwin et al., 2016; Löw et al., 2015; Rösler & Gamer, 2019; Sege et al., 2018). Furthermore, defensive reactions in response to avoidable vs. unavoidable threats recruit different brain regions (Wendt et al., 2017), which might drive these differential HR responses (Roelofs & Dayan, 2022), and could possibly obfuscate potential relations with trait anxiety. Therefore, different situational characteristics might veil the link between trait anxiety and HR responses to threat due to their own inherent associations with trait anxiety, which could explain differences in results across studies. Where bradycardia may be useful for adequate coping in a threat anticipatory state, it may be less useful during circa strike (Roelofs & Dayan, 2022).

Most previous studies on the relation between trait anxiety and associative threat learning used only sympathetic measures (SCR) and showed inconsistent relations with trait anxiety. On the one hand, Sjouwerman and colleagues (2020) did not show SCR threat responses to be linked to trait anxiety, but found that increased trait anxiety was linked to impaired discrimination between threatening and safe stimuli, while Haaker and colleagues (2015) found no discernible link between trait anxiety and ANS responses to threat. While methodological design choices such as reinforcement rate, inclusion of startle probes, or the choice of the conditioned stimuli can have an effect on the threat responses (Lonsdorf et al., 2017; Sjouwerman et al., 2016), it is also possible that the chosen outcome measure plays a vital role in detecting individual differences in threat responses. In the abovementioned studies, threat responses were quantified using sympathetically driven SCRs, a measure that is less sensitive to parasympathetic contributions in the threat response. Our data, in line with previous literature (Stone et al., 2020; Vinkers et al., 2021) suggests that the balance between sympathetic and parasympathetic contribution to threat maybe a better predictor of anxiety related psychopathology.

In conclusion, our results highlight the importance of including parasympathetic as well as sympathetic contributions in the threat response when assessing individual differences in physiological responses to threat. Threat-induced bradycardia and tachycardia each constitute unique indices of learned threat that are differentially linked to individual differences in trait anxiety. Since threat learning processes can yield insights into the aetiology, maintenance, and treatment of anxiety disorders (Mineka & Oehlberg, 2008; VanElzakker et al., 2014), adequate measures of (individual differences in) threat responses are needed in order to best inform clinical practice (Bach et al., 2023). We call for a paradigm shift when assessing individual differences to threat advocating to not classify, nor exclude, participants based on absence of SCR responses, but instead including measures that are sensitive to parasympathetic (as well as sympathetic) influences such as HR. Defensive responses to threat vary across a larger spectrum than sympathetic arousal alone. This is particularly relevant for studies that aim to link threat learning processes to resilience.

## Acknowledgements

KR and LdV were supported by a consolidator grant from the European Research Council (ERC_CoG-2017_772337) awarded to KR.

## References

Adenauer, H., Catani, C., Keil, J., Aichinger, H., & Neuner, F. (2010). Is freezing an adaptive reaction to threat? Evidence from heart rate reactivity to emotional pictures in victims of war and torture. Psychophysiology, 47(2), 315–322. https://doi.org/10.1111/j.1469-8986.2009.00940.x

Ahrens, L. M., Pauli, P., Reif, A., Mühlberger, A., Langs, G., Aalderink, T., & Wieser, M. J. (2016). Fear conditioning and stimulus generalization in patients with social anxiety disorder. Journal of Anxiety Disorders, 44, 36–46. https://doi.org/10.1016/j.janxdis.2016.10.003

Alexandra Kredlow, M., Pineles, S. L., Inslicht, S. S., Marin, M. F., Milad, M. R., Otto, M. W., & Orr, S. P. (2017). Assessment of skin conductance in African American and Non–African American participants in studies of conditioned fear. Psychophysiology, 54(11), 1741–1754. https://doi.org/10.1111/psyp.12909

Bach, D. R., & Melinscak, F. (2020). Psychophysiological modelling and the measurement of fear conditioning. Behaviour Research and Therapy, 127(January), 103576. https://doi.org/10.1016/j.brat.2020.103576

Bach, D. R., Sporrer, J., Abend, R., Beckers, T., Dunsmoor, J. E., Fullana, M., Gamer, M., Gee, D., Hamm, A., Hartley, C., Herringa, R., Jovanovic, T., Kalisch, R., Knight, D., Lissek, S., Lonsdorf, T., Merz, C., Milad, M., Morriss, J., … Schiller, D. (2023). Consensus design of a calibration experiment for human fear conditioning. Neuroscience and Biobehavioral Reviews, 148, 105146. https://doi.org/10.1016/j.neubiorev.2023.105146

Barr, D. J. (2013). Random effects structure for testing interactions in linear mixed-effects models. Frontiers in Psychology, 4(June), 3–4. https://doi.org/10.3389/fpsyg.2013.00328

Battaglia, S., Orsolini, S., Borgomaneri, S., Barbieri, R., Diciotti, S., & di Pellegrino, G. (2022). Characterizing cardiac autonomic dynamics of fear learning in humans. Psychophysiology, May, 1–16. https://doi.org/10.1111/psyp.14122

Bernstein, D. P., Fink, L., Handelsman, L., & Foote, J. (n.d.). Childhood trauma questionnaire. Assessment of family violence: A handbook for researchers and practitioners. 1998.

Bernstein, R. E., Measelle, J. R., Laurent, H. K., Musser, E. D., & Ablow, J. C. (2013). Sticks and Stones May Break My Bones but Words Relate to Adult Physiology Child Abuse Experience and Women’s Sympathetic Nervous System Response while Self-Reporting Trauma. Journal of Aggression, Maltreatment and Trauma, 22(10), 1117–1136. https://doi.org/10.1080/10926771.2013.850138

Blanca, M. J., Arnau, J., López-Montiel, D., Bono, R., & Bendayan, R. (2013). Skewness and kurtosis in real data samples. Methodology, 9(2), 78–84. https://doi.org/10.1027/1614-2241/a000057

Blanchard, D. C., Griebel, G., Pobbe, R., & Blanchard, R. J. (2011). Risk assessment as an evolved threat detection and analysis process. Neuroscience and Biobehavioral Reviews, 35(4), 991–998. https://doi.org/10.1016/j.neubiorev.2010.10.016

Blanchard, R. J., & Blanchard, D. C. (1968). Escape and avoidance responses to a fear eliciting situation. Psychonomic Science, 13(1), 19–20. https://doi.org/10.3758/bf03342387

Blechert, J., Michael, T., Grossman, P., Lajtman, M., & Wilhelm, F. H. (2007). Autonomic and respiratory characteristics of posttraumatic stress disorder and panic disorder. Psychosomatic Medicine, 69(9), 935–943. https://doi.org/10.1097/PSY.0b013e31815a8f6b

Bradley, M. M., Moulder, B., & Lang, P. J. (2005). When good things go bad: The reflex physiology of defense. Psychological Science, 16(6), 468–473.

Brawman-Mintzer, O., & Lydiard, R. B. (1997). Biological Basis of Generalized Anxiety Disorder. Journal of Clinical Psychiatry, 58(suppl 3), 16–25.

Bryant, R. A., Salmon, K., Sinclair, E., & Davidson, P. (2007). Heart rate as a predictor of posttraumatic stress disorder in children. General Hospital Psychiatry, 29(1), 66–68. https://doi.org/10.1016/j.genhosppsych.2006.10.002

Bürkner, P. C. (2018). Advanced Bayesian multilevel modeling with the R package brms. R Journal, 10(1), 395–411. https://doi.org/10.32614/rj-2018-017

Castegnetti, G., Tzovara, A., Staib, M., Paulus, P. C., Hofer, N., & Bach, D. R. (2016). Modeling fear-conditioned bradycardia in humans. Psychophysiology, 53(6), 930–939. https://doi.org/10.1111/psyp.12637

Cohen, D. H., & Randall, D. C. (1984). Classical conditioning of cardiovascular responses. Annual Review of Physiology, *VOL.* 46(c), 187–197. https://doi.org/10.1146/annurev.ph.46.030184.001155

de Echegaray, J., & Moratti, S. (2021). Threat imminence modulates neural gain in attention and motor relevant brain circuits in humans. Psychophysiology, 58(8), 1–16. https://doi.org/10.1111/psyp.13849

de Voogd, L. D., Fernández, G., & Hermans, E. J. (2016). Awake reactivation of emotional memory traces through hippocampal-neocortical interactions. NeuroImage, 134, 563–572. https://doi.org/10.1016/j.neuroimage.2016.04.026

de Voogd, L. D., Hagenberg, E., Zhou, Y. J., de Lange, F. P., & Roelofs, K. (2022). Acute threat enhances perceptual sensitivity without affecting the decision criterion. Scientific Reports, 12(1), 1–11. https://doi.org/10.1038/s41598-022-11664-0

Duits, P., Cath, D. C., Lissek, S., Hox, J. J., Hamm, A. O., Engelhard, I. M., Van Den Hout, M. A., & Baas, J. M. P. (2015). Updated meta-analysis of classical fear conditioning in the anxiety disorders. Depression and Anxiety, 32(4), 239–253. https://doi.org/10.1002/da.22353

Dymond, S., Dunsmoor, J. E., Vervliet, B., Roche, B., & Hermans, D. (2015). Fear Generalization in Humans: Systematic Review and Implications for Anxiety Disorder Research. Behavior Therapy, 46(5), 561–582. https://doi.org/10.1016/j.beth.2014.10.001

Eilam, D. (2005). Die hard: A blend of freezing and fleeing as a dynamic defense - Implications for the control of defensive behavior. Neuroscience and Biobehavioral Reviews, 29(8), 1181–1191. https://doi.org/10.1016/j.neubiorev.2005.03.027

Fanselow, M. S. (1994). Neural organization of the defensive behavior system responsible for fear. Psychonomic Bulletin & Review, 1(4), 429–438. https://doi.org/10.3758/BF03210947

Forsyth, J. P., Daleiden, E. L., & Chorpita, B. F. (2000). Response primacy in fear conditioning: Disentangling the contributions of UCS vs. UCR intensity. Psychological Record, 50(1), 17–33. https://doi.org/10.1007/BF03395340

Fragkaki, I., Roelofs, K., Stins, J., Jongedijk, R. A., & Hagenaars, M. A. (2017). Reduced freezing in posttraumatic stress disorder patients while watching affective pictures. Frontiers in Psychiatry, 8(MAR), 1–9. https://doi.org/10.3389/fpsyt.2017.00039

Gladwin, T. E., Hashemi, M. M., van Ast, V., & Roelofs, K. (2016). Ready and waiting: Freezing as active action preparation under threat. Neuroscience Letters, 619, 182–188. https://doi.org/10.1016/j.neulet.2016.03.027

Green, S. R., Kragel, P. A., Fecteau, M. E., & LaBar, K. S. (2014). Development and validation of an unsupervised scoring system (Autonomate) for skin conductance response analysis. International Journal of Psychophysiology, 91(3), 186–193. https://doi.org/doi:10.1016/j.ijpsycho.2013.10.015

Haaker, J., Lonsdorf, T. B., Schümann, D., Menz, M., Brassen, S., Bunzeck, N., Gamer, M., & Kalisch, R. (2015). Deficient inhibitory processing in trait anxiety: Evidence from context-dependent fear learning, extinction recall and renewal. Biological Psychology, 111, 65–72. https://doi.org/10.1016/j.biopsycho.2015.07.010

Haaker, J., Maren, S., Andreatta, M., Merz, C. J., Richter, J., Richter, S. H., Meir Drexler, S., Lange, M. D., Jüngling, K., Nees, F., Seidenbecher, T., Fullana, M. A., Wotjak, C. T., & Lonsdorf, T. B. (2019). Making translation work: Harmonizing cross-species methodology in the behavioural neuroscience of Pavlovian fear conditioning. Neuroscience and Biobehavioral Reviews, 107(April), 329–345. https://doi.org/10.1016/j.neubiorev.2019.09.020

Hagenaars, M. A., Oitzl, M., & Roelofs, K. (2014). Updating freeze: Aligning animal and human research. Neuroscience and Biobehavioral Reviews, 47, 165–176. https://doi.org/10.1016/j.neubiorev.2014.07.021

Hagenaars, M. A., Roelofs, K., & Stins, J. F. (2014). Human freezing in response to affective films. Anxiety, Stress and Coping, 27(1), 27–37. https://doi.org/10.1080/10615806.2013.809420

Hagenaars, M. A., Stins, J. F., & Roelofs, K. (2012). Aversive life events enhance human freezing responses. In Journal of Experimental Psychology: General (Vol. 141, pp. 98–105). American Psychological Association. https://doi.org/10.1037/a0024211

Hamm, A. O., Greenwald, M. K., Bradley, M. M., & Lang, P. J. (1993). Emotional learning, hedonic change, and the startle probe. In Journal of Abnormal Psychology (Vol. 102, Issue 3, pp. 453–465). https://doi.org/10.1037//0021-843x.102.3.453

Hashemi, M. M., Gladwin, T. E., de Valk, N. M., Zhang, W., Kaldewaij, R., van Ast, V., Koch, S. B. J., Klumpers, F., & Roelofs, K. (2019). Neural Dynamics of Shooting Decisions and the Switch from Freeze to Fight. Scientific Reports, 9(1), 1–10. https://doi.org/10.1038/s41598-019-40917-8

Hashemi, M. M., Zhang, W., Kaldewaij, R., Koch, S. B. J., Smit, A., Figner, B., Jonker, R., Klumpers, F., & Roelofs, K. (2021). Human defensive freezing: Associations with hair cortisol and trait anxiety. Psychoneuroendocrinology, 133(September 2020), 105417. https://doi.org/10.1016/j.psyneuen.2021.105417

Held, L. K., Vink, J. M., Vitaro, F., Brendgen, M., Dionne, G., Provost, L., Boivin, M., Ouellet-Morin, I., & Roelofs, K. (2022). The gene environment aetiology of freezing and its relationship with internalizing symptoms during adolescence. EBioMedicine, 81, 104094. https://doi.org/10.1016/j.ebiom.2022.104094

Hermans, E. J., Henckens, M. J. A. G., Roelofs, K., & Fernández, G. (2013). Fear bradycardia and activation of the human periaqueductal grey. NeuroImage, 66, 278–287. https://doi.org/10.1016/j.neuroimage.2012.10.063

Hodes, R. L., Cook, E. W., & Lang, P. J. (1985). Individual Differences in Autonomic Response: Conditioned Association or Conditioned Fear? Psychophysiology, 22(5), 545–560.

Hulsman, A. M., Kaldewaij, R., Hashemi, M. M., Zhang, W., Koch, S. B. J., Figner, B., Roelofs, K., & Klumpers, F. (2021). Individual differences in costly fearful avoidance and the relation to psychophysiology. Behaviour Research and Therapy, 137(June 2020), 103788. https://doi.org/10.1016/j.brat.2020.103788

Iwata, J., & LeDoux, J. E. (1988). Dissociation of Associative and Nonassociative Concommitants of Classical Fear Conditioning in the Freely Behaving Rat. Behavioral Neuroscience, 102(1), 66–76. https://doi.org/10.1037/0735-7044.102.1.66

Jaswetz, L., de Voogd, L. D., Becker, E. S., & Roelofs, K. (2022). No evidence for disruption of reconsolidation of conditioned threat memories with a cognitively demanding intervention. Scientific Reports, 12(1), 1–12. https://doi.org/10.1038/s41598-022-10184-1

Keay, K. A., & Bandler, R. (2001). Parallel circuits mediating distinct emotional coping reactions to different types of stress. Neuroscience and Biobehavioral Reviews, 25(7–8), 669–678. https://doi.org/10.1016/S0149-7634(01)00049-5

Klaassen, F. H., Held, L., Figner, B., O’Reilly, J. X., Klumpers, F., de Voogd, L. D., & Roelofs, K. (2021). Defensive freezing and its relation to approach–avoidance decision-making under threat. Scientific Reports, 11(1), 1–12. https://doi.org/10.1038/s41598-021-90968-z

Klorman, R., & Ryan, R. M. (1980). Heart Rate, Contingent Negative Variation, and Evoked Potentials during Anticipation of Affective Stimulation. Psychophysiology, 17(6), 513–523.

Klumpers, F., Van Gerven, J. M., Prinssen, E. P. M., Niklson Roche, I., Roesch, F., Riedel, W. J., Kenemans, J. L., & Baas, J. M. P. (2010). Method development studies for repeatedly measuring anxiolytic drug effects in healthy humans. Journal of Psychopharmacology, 24(5), 657–666. https://doi.org/10.1177/0269881109103115

Knowles, K. A., & Olatunji, B. O. (2020). Specificity of trait anxiety in anxiety and depression: Meta-analysis of the State-Trait Anxiety Inventory. Clinical Psychology Review, 82(September), 101928. https://doi.org/10.1016/j.cpr.2020.101928

Lipov, E. (2013). Post Traumatic Stress Disorder (PTSD) as an Over Activation of Sympathetic Nervous System: An Alternative View. Trauma & Treatment, 03(01), 1–3. https://doi.org/10.4172/2167-1222.1000181

Livermore, J. J. A., Klaassen, F. H., Bramson, B., Hulsman, A. M., Meijer, S. W., Held, L., Klumpers, F., de Voogd, L. D., & Roelofs, K. (2021). Approach-Avoidance Decisions Under Threat: The Role of Autonomic Psychophysiological States. Frontiers in Neuroscience, 15(March), 1–12. https://doi.org/10.3389/fnins.2021.621517

Lojowska, M., Gladwin, T. E., & Roelofs, K. (2015). Supplemental Material for Freezing Promotes Perception of Coarse Visual Features. Journal of Experimental Psychology: General, 144(6), 1080–1088. https://doi.org/10.1037/xge0000117.supp

Lojowska, M., Ling, S., Roelofs, K., & Hermans, E. J. (2018). Visuocortical changes during a freezing-like state in humans. NeuroImage, 179(June), 313–325. https://doi.org/10.1016/j.neuroimage.2018.06.013

Lonsdorf, T. B., Klingelhöfer-Jens, M., Andreatta, M., Beckers, T., Chalkia, A., Gerlicher, A., Jentsch, V. L., Drexler, S. M., Mertens, G., Richter, J., Sjouwerman, R., Wendt, J., & Merz, C. J. (2019). Navigating the garden of forking paths for data exclusions in fear conditioning research. ELife, 8, 1–36. https://doi.org/10.7554/eLife.52465

Lonsdorf, T. B., Menz, M. M., Andreatta, M., Fullana, M. A., Golkar, A., Haaker, J., Heitland, I., Hermann, A., Kuhn, M., Kruse, O., Meir Drexler, S., Meulders, A., Nees, F., Pittig, A., Richter, J., Römer, S., Shiban, Y., Schmitz, A., Straube, B., … Merz, C. J. (2017). Don’t fear ‘fear conditioning’: Methodological considerations for the design and analysis of studies on human fear acquisition, extinction, and return of fear. Neuroscience and Biobehavioral Reviews, 77, 247–285. https://doi.org/10.1016/j.neubiorev.2017.02.026

Löw, A., Lang, P. J., Smith, J. C., & Bradley, M. M. (2008). Both predator and prey: Emotional arousal in threat and reward. Psychological Science, 19(9), 865–873. https://doi.org/10.1111/j.1467-9280.2008.02170.x

Löw, A., Weymar, M., & Hamm, A. O. (2015). When Threat Is Near, Get Out of Here: Dynamics of Defensive Behavior During Freezing and Active Avoidance. Psychological Science, 26(11), 1706– 1716. https://doi.org/10.1177/0956797615597332

Ly, V., Huys, Q. J. M., Stins, J. F., Roelofs, K., & Cools, R. (2014). Individual differences in bodily freezing predict emotional biases in decision making. Frontiers in Behavioral Neuroscience, 8(JULY), 1–9. https://doi.org/10.3389/fnbeh.2014.00237

Marin, M. F., Zsido, R. G., Song, H., Lasko, N. B., Killgore, W. D. S., Rauch, S. L., Simon, N. M., & Milad, M. R. (2017). Skin conductance responses and neural activations during fear conditioning and extinction recall across anxiety disorders. JAMA Psychiatry, 74(6), 622–631. https://doi.org/10.1001/jamapsychiatry.2017.0329

MATLAB (No. 2018b). (2018). The MathWorks Inc.

Minassian, A., Maihofer, A. X., Baker, D. G., Nievergelt, C. M., Geyer, M. A., Risbrough, V. B., Hauger, R. L., Huang, M., Murphy, J. A., Naviaux, R. K., Yurgil, K., Patel, A., De La Rosa, A., & Gorman, P. (2015). Association of predeployment heart rate variability with risk of postdeployment posttraumatic stress disorder in active-duty marines. JAMA Psychiatry, 72(10), 979–986. https://doi.org/10.1001/jamapsychiatry.2015.0922

Mineka, S., & Oehlberg, K. (2008). The relevance of recent developments in classical conditioning to understanding the etiology and maintenance of anxiety disorders. Acta Psychologica, 127(3), 567–580. https://doi.org/10.1016/j.actpsy.2007.11.007

Mobbs, D. (2018). The ethological deconstruction of fear(s). Current Opinion in Behavioral Sciences, 24, 32–37. https://doi.org/10.1016/j.cobeha.2018.02.008

Mobbs, D., Headley, D. B., Ding, W., & Dayan, P. (2020). Space, Time, and Fear: Survival Computations along Defensive Circuits. Trends in Cognitive Sciences, 24(3), 228–241. https://doi.org/10.1016/j.tics.2019.12.016

Moratti, S., & Keil, A. (2005). Cortical activation during Pavlovian fear conditioning depends on heart rate response patterns: An MEG study. Cognitive Brain Research, 25(2), 459–471. https://doi.org/10.1016/j.cogbrainres.2005.07.006

Morris, M. C., & Rao, U. (2013). Psychobiology of PTSD in the acute aftermath of trauma: Integrating research on coping, HPA function and sympathetic nervous system activity. Asian Journal of Psychiatry, 6(1), 3–21. https://doi.org/10.1016/j.ajp.2012.07.012

Niermann, H. C. M., Figner, B., & Roelofs, K. (2017). Individual differences in defensive stress-responses: the potential relevance for psychopathology. Current Opinion in Behavioral Sciences, 14, 94–101. https://doi.org/10.1016/j.cobeha.2017.01.002

Niermann, H. C. M., Figner, B., Tyborowska, A., Cillessen, A. H. N., & Roelofs, K. (2018). Investigation of the stability of human freezing-like responses to social threat from mid to late adolescence. Frontiers in Behavioral Neuroscience, 12(May), 1–9. https://doi.org/10.3389/fnbeh.2018.00097

Noordewier, M. K., Scheepers, D. T., & Hilbert, L. P. (2020). Freezing in response to social threat: a replication. Psychological Research, 84(7), 1890–1896. https://doi.org/10.1007/s00426-019-01203-4

Panitz, C., Hermann, C., & Mueller, E. M. (2015). Conditioned and extinguished fear modulate functional corticocardiac coupling in humans. Psychophysiology, 52(10), 1351–1360. https://doi.org/10.1111/psyp.12498

Pittig, A., Boschet, J. M., Glück, V. M., & Schneider, K. (2021). Elevated costly avoidance in anxiety disorders: Patients show little downregulation of acquired avoidance in face of competing rewards for approach. Depression and Anxiety, 38(3), 361–371. https://doi.org/10.1002/da.23119

Pittig, A., & Scherbaum, S. (2020). Costly avoidance in anxious individuals: Elevated threat avoidance in anxious individuals under high, but not low competing rewards. Journal of Behavior Therapy and Experimental Psychiatry, 66(December 2018), 101524. https://doi.org/10.1016/j.jbtep.2019.101524

Pittig, A., Treanor, M., LeBeau, R. T., & Craske, M. G. (2018). The role of associative fear and avoidance learning in anxiety disorders: Gaps and directions for future research. Neuroscience and Biobehavioral Reviews, 88(October 2017), 117–140. https://doi.org/10.1016/j.neubiorev.2018.03.015

Plunkett, R. P. (1979). Effect of hippocampal lesions on heart-rate during classical fear conditioning. Physiology and Behavior, 23(3), 433–437. https://doi.org/10.1016/0031-9384(79)90039-8

Pole, N. (2007). The Psychophysiology of Posttraumatic Stress Disorder: A Meta-Analysis. Psychological Bulletin, 133(5), 725–746. https://doi.org/10.1037/0033-2909.133.5.725

Qi, S., Hassabis, D., Sun, J., Guo, F., Daw, N., & Mobbs, D. (2018). How cognitive and reactive fear circuits optimize escape decisions in humans. Proceedings of the National Academy of Sciences of the United States of America, 115(12), 3186–3191. https://doi.org/10.1073/pnas.1712314115

R Core Team. (2020). *R: A language and environment for statistical computing*. https://www.r-project.org/

Roelofs, K. (2017). Freeze for action: Neurobiological mechanisms in animal and human freezing. Philosophical Transactions of the Royal Society B: Biological Sciences, 372(1718). https://doi.org/10.1098/rstb.2016.0206

Roelofs, K., & Dayan, P. (2022). Freezing revisited: coordinated autonomic and central optimization of threat coping. Nature Reviews Neuroscience, 23(9), 568–580. https://doi.org/10.1038/s41583-022-00608-2

Roelofs, K., Hagenaars, M. A., & Stins, J. (2010). Facing freeze: Social threat induces bodily freeze in humans. Psychological Science, 21(11), 1575–1581. https://doi.org/10.1177/0956797610384746

Rösler, L., & Gamer, M. (2019). Freezing of gaze during action preparation under threat imminence. Scientific Reports, 9(1), 1–9. https://doi.org/10.1038/s41598-019-53683-4

Sandin, B., & Chorot, P. (1989). The incubation theory of fear/anxiety: Experimental investigation in a human laboratory model of Pavlovian conditioning. Behaviour Research and Therapy, 27(1), 9–18. https://doi.org/10.1016/0005-7967(89)90114-9

Sege, C. T., Bradley, M. M., & Lang, P. J. (2018). Avoidance and escape: Defensive reactivity and trait anxiety. Behaviour Research and Therapy, 104(November 2017), 62–68. https://doi.org/10.1016/j.brat.2018.03.002

Sep, M. S. C., Steenmeijer, A., & Kennis, M. (2019). The relation between anxious personality traits and fear generalization in healthy subjects: A systematic review and meta-analysis. Neuroscience and Biobehavioral Reviews, 107(September), 320–328. https://doi.org/10.1016/j.neubiorev.2019.09.029

Sevenster, D., Hamm, A., Beckers, T., & Kindt, M. (2015). Heart rate pattern and resting heart rate variability mediate individual differences in contextual anxiety and conditioned responses. International Journal of Psychophysiology, 98(3), 567–576. https://doi.org/10.1016/j.ijpsycho.2015.09.004

Sjouwerman, R., Niehaus, J., Kuhn, M., & Lonsdorf, T. B. (2016). Don’t startle me—Interference of startle probe presentations and intermittent ratings with fear acquisition. Psychophysiology, 53(12), 1889–1899. https://doi.org/10.1111/psyp.12761

Sjouwerman, R., Scharfenort, R., & Lonsdorf, T. B. (2020). Individual differences in fear acquisition: multivariate analyses of different emotional negativity scales, physiological responding, subjective measures, and neural activation. Scientific Reports, 10(1), 1–20. https://doi.org/10.1038/s41598-020-72007-5

Spielberger, C. D. (1983). *State-trait anxiety inventory for adults*.

Stone, L. B., McCormack, C. C., & Bylsma, L. M. (2020). Cross system autonomic balance and regulation: Associations with depression and anxiety symptoms. Psychophysiology, 57(10), 1–10. https://doi.org/10.1111/psyp.13636

Szeska, C., Richter, J., Wendt, J., Weymar, M., & Hamm, A. O. (2021). Attentive immobility in the face of inevitable distal threat—Startle potentiation and fear bradycardia as an index of emotion and attention. Psychophysiology, 58(6), 1–17. https://doi.org/10.1111/psyp.13812

Tavakol, M., & Dennick, R. (2011). Making sense of Cronbach’s alpha. In International journal of medical education (Vol. 2, pp. 53–55). https://doi.org/10.5116/ijme.4dfb.8dfd

Trott, J. M., Hoffman, A. N., Zhuravka, I., & Fanselow, M. S. (2022). Conditional and Unconditional Components of Aversively Motivated Freezing, Flight and Darting in Mice. ELife, 11, 1–25. https://doi.org/10.7554/eLife.75663

van Ast, V. A., Klumpers, F., Grasman, R. P. P. P., Krypotos, A. M., & Roelofs, K. (2022). Postural freezing relates to startle potentiation in a human fear-conditioning paradigm. Psychophysiology, 59(4). https://doi.org/10.1111/psyp.13983

Van Diest, I., Bradley, M. M., Guerra, P., Van den Bergh, O., & Lang, P. J. (2009). Fear-conditioned respiration and its association to cardiac reactivity. Biological Psychology, 80(2), 212–217. https://doi.org/10.1016/j.biopsycho.2008.09.006

VanElzakker, M. B., Kathryn Dahlgren, M., Caroline Davis, F., Dubois, S., & Shin, L. M. (2014). From Pavlov to PTSD: The extinction of conditioned fear in rodents, humans, and anxiety disorders. Neurobiology of Learning and Memory, 113, 3–18. https://doi.org/10.1016/j.nlm.2013.11.014

Vinkers, C. H., Kuzminskaite, E., Lamers, F., Giltay, E. J., & Penninx, B. W. J. H. (2021). An integrated approach to understand biological stress system dysregulation across depressive and anxiety disorders. Journal of Affective Disorders, 283(January), 139–146. https://doi.org/10.1016/j.jad.2021.01.051

Walker, P., & Carrive, P. (2003). Role of ventrolateral periaqueductal gray neurons in the behavioral and cardiovascular responses to contextual conditioned fear and poststress recovery. Neuroscience, 116(3), 897–912. https://doi.org/10.1016/S0306-4522(02)00744-3

Waxenbaum, J. A., Reddy, V., & Varacallo, M. (2021). Anatomy, autonomic nervous system. StatPearls Publishing.

Wendt, J., Löw, A., Weymar, M., Lotze, M., & Hamm, A. O. (2017). Active avoidance and attentive freezing in the face of approaching threat. NeuroImage, 158(December 2016), 196–204. https://doi.org/10.1016/j.neuroimage.2017.06.054

Wickramasuriya, D. S., & Faghih, R. T. (2020). A mixed filter algorithm for sympathetic arousal tracking from skin conductance and heart rate measurements in Pavlovian fear conditioning. In PLoS ONE (Vol. 15, Issue 4). https://doi.org/10.1371/journal.pone.0231659

